# Retinoic acid signaling mediates peripheral cone photoreceptor survival in a mouse model of retina degeneration

**DOI:** 10.1101/2021.12.14.472645

**Authors:** Ryoji Amamoto, Grace K. Wallick, Constance L. Cepko

## Abstract

Retinitis Pigmentosa (RP) is a wide array of progressive, debilitating visual disorders caused by mutations in a diverse set of genes. In both human patients and mouse models of RP, rod photoreceptor dysfunction leads to loss of night vision, and is followed by secondary cone photoreceptor dysfunction and degeneration, leading to loss of daylight color vision. A strategy to prevent secondary cone death could provide a generalized RP therapy to preserve daylight color vision regardless of the underlying mutation. In mouse models of RP, cones in the far peripheral retina survive long-term, despite complete rod loss. The mechanism for such peripheral cone survival had not been explored. Here, we found that active retinoic acid (RA) signaling in peripheral Muller glia is both sufficient and necessary for the extended cone survival. RA depletion by conditional knockout of RA synthesis enzymes, or overexpression of an RA degradation enzyme, abrogated peripheral cone survival. Conversely, constitutive activation of RA signaling in the central retina promoted long-term cone survival. These results indicate that RA signaling mediates the prolonged peripheral cone survival in the rd1 mouse model of retinal degeneration, and provide a basis for a generic strategy for cone survival in the many diseases that lead to loss of cone-mediated vision.

## INTRODUCTION

Retinitis Pigmentosa (RP) is a disease of the retina that affects 1 in 4000 people worldwide^1^. It is characterized by an initial loss of night vision, followed by loss of color daylight vision. These symptoms are caused by the progressive degeneration of photoreceptors, rods and cones, which are necessary for achromatic night vision and daylight color vision, respectively. Although the disease progression is variable, a typical patient with RP loses night vision as an adolescent and subsequently loses central daylight vision by the age of 60^1^. Although early diagnosis is possible by procedures such as full-field retinal electroretinogram and dark adaptometry, there is no effective therapy for the vast majority of patients. Although gene therapy via gene augmentation has proven to be effective in humans^2-5^, a one-by-one genetic augmentation approach is currently not feasible given that 69 causative genes for RP have been identified (RetNet). However, a generic therapy aimed at preserving cone-mediated vision may be possible. Many RP disease genes are expressed solely in rods, causing rod dysfunction and death^6^. Cones then suffer from bystander effects, leading to poor daylight and color vision, and in many cases, complete blindness.

The mechanisms underlying the non-cell autonomous, rod-dependent cone degeneration in RP are not completely understood. Previous studies have suggested different mechanisms to explain this secondary cone death, including toxicity from the degenerating rods, loss of rod-derived survival factors, oxidative damage, immune cell activation, necroptosis, and altered metabolism^7-18^. Ameliorating one or several of these mechanisms might then provide a generalized RP therapy to preserve daylight color vision. Intriguingly, we and others have noted that, in animal models of RP, the cones in the far periphery survive^6,19-27^. The basis of this extended cone survival had not been addressed, though one hypothesis is that light exposure, which can induce photoreceptor degeneration^28^, is reduced in the far periphery. However, while dark rearing can slow photoreceptor degeneration, particularly in albino RP animal models, it does not rescue the degeneration phenotype^29-33^. Additionally, cone survival in the far periphery is confined to an area demarcated by a sharp boundary, as opposed to a gradient, which would be expected if light exposure was the sole explanation. These results indicate that cone survival in the far periphery of the retina is not solely mediated by different levels of light exposure. Therefore, we asked whether there are molecular determinants that regulate cone survival in the far periphery, and if so, whether such determinants are sufficient to promote cone survival in the central retina.

We found that the regions of cone survival in the peripheral retina overlap with areas of active retinoic acid (RA) signaling. RA signaling regulates a wide range of activities such as cellular differentiation and survival in various organs including the central nervous system^34,35^, but has not been reported to play a role in photoreceptor survival. In canonical RA signaling, RA is synthesized by Aldh1a1, Aldh1a2, or Aldh1a3, and it binds to RA receptors (RAR), which activate transcription of RA target genes via binding to genomic RA Response Elements (RARE)^34^. During development of the mouse eye, *Aldh1a1* and *Aldh1a3* are expressed in the dorsal and ventral retinal periphery, respectively, with the expression of dorsal *Aldh1a1* remaining into adulthood^36-38^. Deletion of the RA synthesis enzymes or the RA receptor in mice has revealed a role for paracrine RA signaling in the perioptic mesenchyme and choroidal vasculature, but not in retina patterning^39-43^. Conditional knock-out of all three synthesis enzymes in adult mice resulted in corneal thinning^44^. Through loss-of-function experiments, we found that RA produced from Muller glia (MG), the major non-neuronal cell type of the retina, is necessary for peripheral cone survival. In addition, we found that constitutive RA signaling activation in MG is sufficient to promote cone survival in the more central retina. Furthermore, we found that *ALDH1A1* expression is enriched in the human peripheral retina, suggesting that this mechanism of cone survival may play a role in human patients with RP.

## RESULTS

### Cone photoreceptors survive in the peripheral retina where RA signaling is active

The rd1 mouse is a rapid photoreceptor degeneration model. It harbors a viral insertion and a point mutation in the Pde6b gene^45^, which is expressed only in rods. The rod death is considered a direct result of Pde6b loss-of-function **(Figure 1A)**. Degeneration of rod photoreceptors starts at approximately P11, and rapidly progresses, such that most rods degenerate by P20. As is the case with most RP cases in animal models and humans, cone photoreceptors then begin to lose function and degenerate^11,27^ **(Figure 1A)**. From the perspective of a retinal flatmount, both rods and cones degenerate in a center-to-periphery pattern. However, we and others^6,19-27^ have noted that cones in the peripheral retina survive longer, although neither the extent nor the mechanism(s) of the extended peripheral cone survival had been explored.

**Figure 1:**
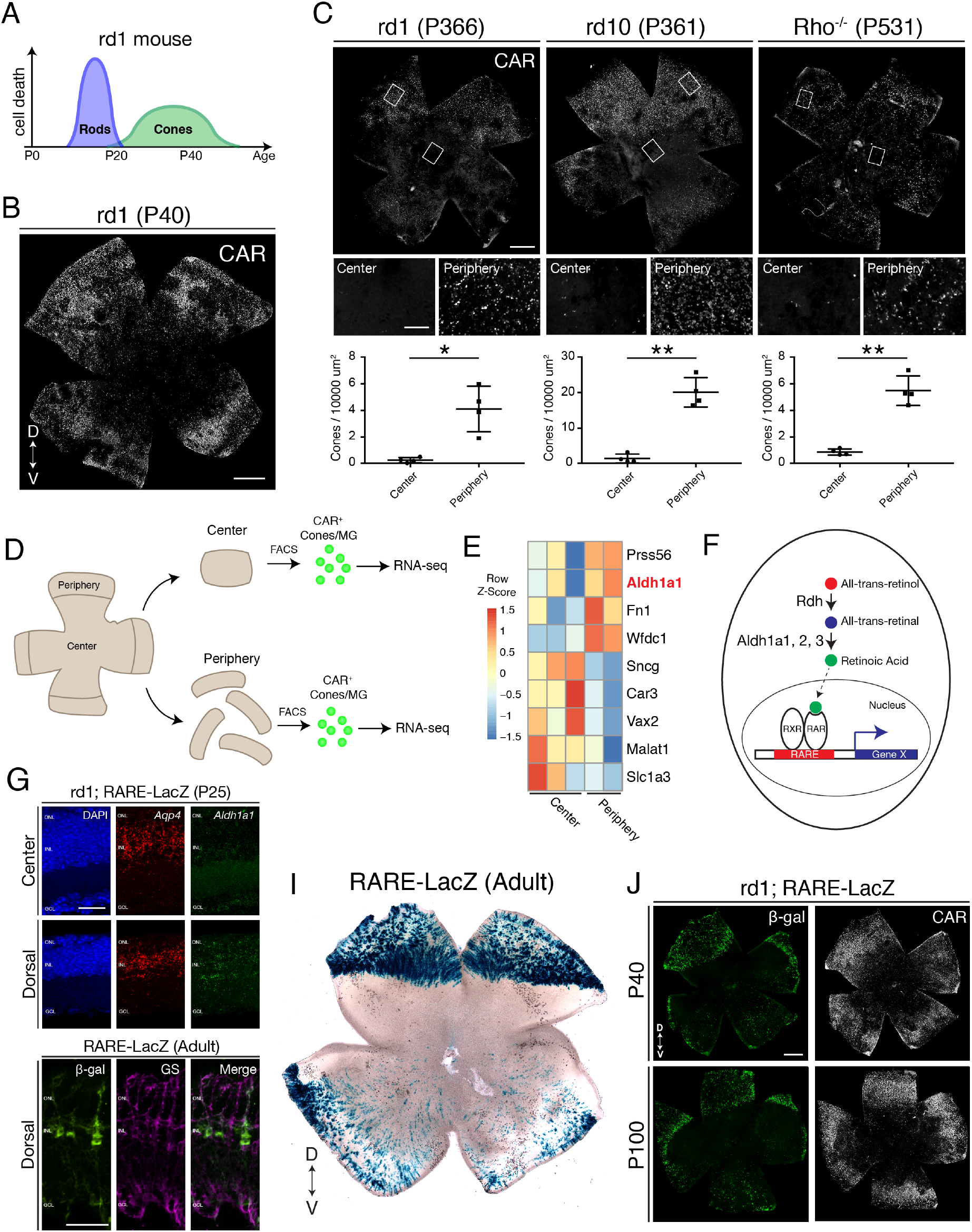
Cones in the rd1 mouse model survive long-term in regions with active RA signaling. (**A**) Schematic of the timeline of rod and cone photoreceptor death in the rd1 mouse model. (**B**) rd1 retinal flatmount IHC against Cone Arrestin (CAR) at P40. (**C**) Retinal flatmount IHC against CAR for rd1, rd10, and Rho^-/-^mouse models at P366, P361, and P531, respectively (top images). Center and peripheral insets used for cone quantification (middle images). Number of CAR^+^ cones/10,000 *μ*m2 (n=4 each) in the center and periphery for rd1 (Student’s two-tailed T test, *p*=0.0206), rd10 (Student’s two-tailed T test, *p*=0.0036), and Rho^-/-^(Student’s two-tailed T test, *p*=0.0041). (**D**) Schematic of the strategy for cone-specific bulk RNA sequencing. Central and peripheral CD1 (non-degenerating strain) retinal tissues were collected, and CAR^+^ cones (and contaminating MG) were FACS purified for downstream RNA sequencing. (**E**) A heatmap representing relative expression levels of differentially expressed genes (adjusted *p*-value*<*0.05) between central and peripheral retina. (**F**) Simplified schematic of the RA signaling pathway. (**G**) SABER smFISH against *Aqp4*, a pan MG marker, and *Aldh1a1* in P25 rd1; RARE-LacZ dorsal and central retinal sections. (**H**) IHC against B-gal and GS, a MG marker, in adult RARE-LacZ retinal sections. (**I**) LacZ staining on adult RARE-LacZ flatmount. (**J**) IHC against B-gal and CAR in P40 (top row) and P100 (bottom row) rd1; RARE-LacZ flatmounts. D, Dorsal; V, Ventral; ONL, Outer Nuclear Layer; INL, Inner Nuclear Layer; GCL, Ganglion Cell Layer. Scale bars; 500 μm (B, C, J), 50 μm (G, H). All results are expressed as the mean*±*SD. *p*<*0.05, **p*<*0.01.

To determine whether such peripheral cone survival persists long-term, we performed immunohistochemical (IHC) analysis of retinal flatmounts, which allows imaging of all cones in the retina. In agreement with previous studies, we found that Cone Arrestin^+^ (CAR; also known as Arr3) cones were enriched in the peripheral retina at P40, with a preference towards the dorsal region **(Figure 1B)**. CAR^+^ cones in the dorsal periphery persisted at least up to P366, and interestingly, a nearly straight boundary demarcated the zone of survival **(Figure 1C, left panel)**. Quantification of the number of cones in sampled peripheral and central regions showed that the peripheral retina contained significantly more cones compared to the central retina **(Figure 1C)**. Such long-term peripheral cone survival was not restricted to the rd1 mouse model. The rd10 mouse model harbors a different mutation in the Pde6b gene, and the rate of photoreceptor degeneration is slower. Similarly, photoreceptor degeneration proceeds more slowly in the Rho^-/-^ mouse model^46^. In both rd10 and Rho^-/-^ mouse models, cones survived long-term in the peripheral retina, persisting at least up to P366 and P531, respectively **(Figure 1C)**. These results indicate that cones in the peripheral retina, especially dorsally, survive long-term in multiple models of rod-cone degeneration.

Next, we sought to understand the molecular mechanism that regulates cone survival in the periphery. First, we asked whether the peripheral cones were surviving because peripheral rods survived long-term. Previous ultrastructural examination of the rd1 retina revealed complete rod loss by P36^19^. To corroborate, we performed single molecule fluorescent *in situ* hybridization (smFISH) for *Nrl*, a marker of rods, and *Arr3*, a marker of cones, in dorsal rd10 retinal sections at P366. While *Arr3*^+^ cones were sparsely detected in the ONL, no *Nrl* signal was found **(Figure 1 – figure supplement 1)**. These results suggest that cones survive in the peripheral retina despite complete rod loss.

We reasoned that the peripheral cones might maintain an intrinsic transcriptional program that promotes cell survival and/or inhibits cell death. Central and peripheral cones from WT mouse retinas were thus profiled to determine if there were transcriptional differences that might underlie such intrinsic differences. To this end, an antibody-based FACS isolation strategy that we had previously developed for both fresh and frozen brain/retina tissue samples was used^47,48^ **(Figure 1D)**. Using the CAR antibody and FACS, a population of CAR^+^ cells was easily identified and isolated, although the percentage of CAR^+^ cells was higher than expected (Expected: 3%, Observed: 15%). Upon performing ddPCR on the extracted RNA, the FACS-isolated population was found to contain *Arr3*^+^/*Rxrg*^+^ cones and *Glul*^+^ MG, but not rods **(Figure 1 – figure supplement 2)**. Despite the contamination, SMART-Seq v4 cDNA libraries were generated and sequenced on NextSeq 500. The central and peripheral samples were analyzed for differential expression (DE). Among the DE genes, we found that Aldh1a1 was highly enriched in the peripheral samples **(Figure 1E)**, matching the previous data that it is expressed in dorsal MG^37^. Functionally, Aldh1a1, Aldh1a2, and Aldh1a3 catalyze the synthesis of RA (RA), a metabolite that, when bound to RA receptor (RAR), drives transcription of RA target genes from genomic RA response elements (RAREs)^34^ **(Figure 1F)**. To validate the expression pattern of Aldh1a1, we performed smFISH for *Aqp4*, a pan MG marker, and *Aldh1a1*, in P25 rd1 retinal sections. In the central retina, *Aldh1a1* signal was detected only at low levels. However, in the dorsal retina, *Aldh1a1* puncta were distributed throughout the radial dimension of the retina, suggestive of an expression pattern in MG, as MG processes exhibit this widespread pattern **(Figure 1G)**. The expression pattern of Aldh1a1 was also validated at the protein level **(Figure 1 – figure supplement 3)**.

A mouse strain, RARE-LacZ, can be used to identify transcriptional activation of the RA signaling pathway at a cellular level^49^. The read-out can be via IHC of the beta-galactosidase (B-gal) protein or chromogenic detection of its activity using X-gal. In sections, B-gal was mostly localized to cell bodies in the INL and processes spanning the retinal section, reminiscent of MG morphology **(Figure 1H)**. Accordingly, B-gal staining overlapped with that of Glutamine Synthetase, an MG marker, indicating that most cells with activated RA signaling were MG. B-gal was not expressed in any other cell type except ChAT^+^ amacrine cells in the ganglion cell layer **(Figure 1 – figure supplement 4)**. Of note, regions of Aldh1a1 expression, and transcription from RA signaling, as indicated by B-gal expression, closely overlapped. This suggests that the RA synthesized by peripheral MG binds to receptors locally, and that the RA does not diffuse long distances **(Figure 1 – figure supplement 3)**. In retinal flatmounts, the strongest B-gal activity was observed in the dorsal periphery, demarcated with a clear border, and weaker activity was found in the ventral periphery, as previously described^50^ **(Figure 1I)**. To determine if this pattern of expression occurred in a retinal degeneration strain, the RARE-LacZ mice were crossed with rd1 mice. In rd1; RARE-lacZ mice, the spatial pattern of B-gal expression closely matched that of CAR^+^ cone survival in rd1 mice, at both P40 and P100 **(Figure 1J)**.

Taken together, these results indicate that cones in the rd1 mouse model survive long-term in regions which exhibit signaling via the RA receptor. These regions express the RA synthesizing enzyme, Aldh1a1, in MG, the cell type that exhibits active RA signaling.

### RA is necessary for peripheral cone survival

To determine whether RA is necessary for peripheral cone survival in a retinal degeneration strain, mice deficient in synthesis of RA in a retinal degeneration background were constructed. The rd1; RARE-LacZ mice were crossed to CAG-CreER and the Aldh1a1^fl/fl^; Aldh1a2^fl/fl^, Aldh1a3^fl/fl^ mouse lines to generate a rd1; CAG-CreER; Aldh1a1^fl/fl^; Aldh1a2^fl/fl^; Aldh1a3^fl/fl^ mouse line (henceforth referred to as Aldh Flox)^44^ **(Figure 2A)**. CAG-CreER was chosen as the Cre driver line to control the developmental stage at which the RA synthesis enzymes were knocked out, as RA is known to be a critical regulator of the development of many organ systems^34^. The Aldh1a2^fl/fl^ and Aldh1a3^fl/fl^ alleles were included, despite the low expression levels of these genes in the adult retina, to minimize the potential effect of gene compensation. A single dose of tamoxifen was injected intraperitoneally at P10, a time point at which genesis of all retinal cell types is finished, and the retinas were harvested at P60 **(Figure 2A)**. As assessed by ddPCR of the retina, the relative expression level of *Aldh1a1* was significantly reduced in retinas with CAG-CreER, compared to control littermates without CAG-CreER (mean *±* SD: -CreER: 0.01203 *±* 0.005; +CreER: 0.0003 *±* 0.0002) **(Figure 2B)**. In P60 Aldh flox retinal flatmounts without CAG-CreER, IHC for CAR showed abundant cone survival in the periphery. In contrast, those with CAG-CreER appeared to have fewer cones in the periphery **(Figure 2C, D)**. Accordingly, quantification of CAR^+^ cones in sampled dorsal regions showed a significant reduction in the number of cones (mean *±* SD: -CreER: 40.941*±* 3.51; +CreER: 23.91*±* 2.74), suggesting that RA is necessary for peripheral cone survival **(Figure 2E)**.

**Figure 2:**
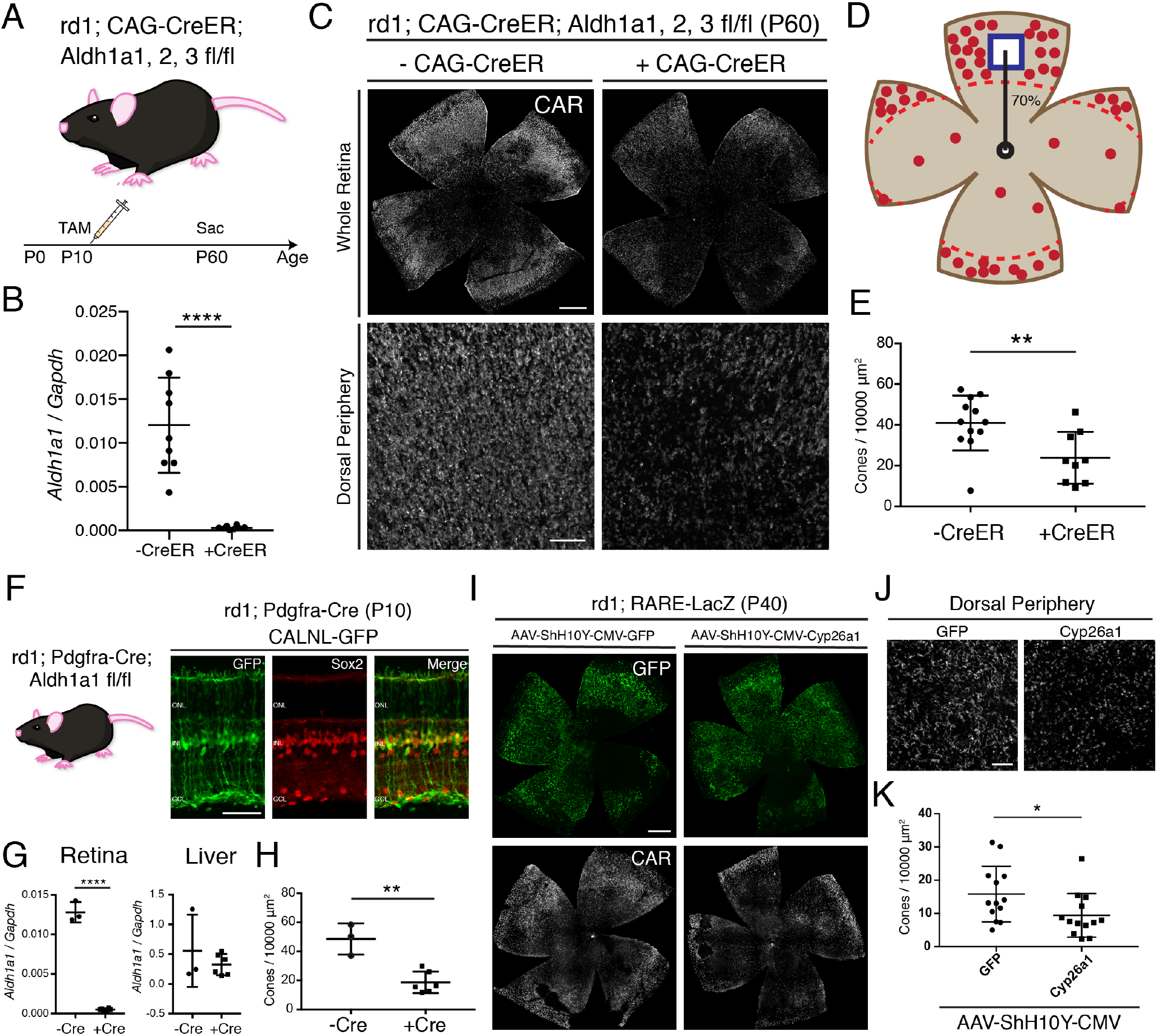
RA is necessary for peripheral cone survival. (**A**) Schematic of the experimental design. Aldh Flox mice were injected with tamoxifen to create a conditional knockout (cKO) of the RA synthetic enzymes at P10, and the retinas were harvested at P60. (**B**) Relative expression level of *Aldh1a1* in retinas with or without CAG-CreER (-Cre: n=9, +Cre: n=8, Student’s two-tailed T test, p*<*0.0001). (**C**) IHC against CAR in P60 Aldh Flox flatmounts showing the entire retina (top row) and the dorsal periphery (bottom row) with or without CAG-CreER. (**D**) Schematic of the area of cone quantification in the dorsal retina. Red dots represent surviving cones, and the blue box is the area of quantification. (**E**) Quantification of CAR^+^ cones in the sampled area of Aldh Flox retinas with or without CAG-CreER (-Cre: n=12, +Cre: n=9, Student’s two-tailed T test, *p*=0.0086). (**F**) Validation of MG-specific Aldh1a1 cKO mouse line. IHC against GFP and Sox2 in P10 rd1; Pdgfra-Cre retinas electroporated with a Cre-dependent plasmid, CALNL-GFP. (**G**) Relative expression level of *Aldh1a1* in retinas and liver with or without Pdgfra-Cre (Retina: -Cre: n=3, +Cre: n=6, Student’s two-tailed T test, p*<*0.0001; Liver: -Cre: n=3, +Cre: n=6, Student’s two-tailed T test, p=0.3893). (**H**) Quantification of CAR^+^ cones in the sampled area of Aldh Flox retinas with or without Pdgfra-Cre (-Cre: n=3, +Cre: n=6, Student’s two-tailed T test, *p*=0.0016). (**I**) IHC against GFP (top row) and CAR (bottom row) in P40 rd1; RARE-LacZ flatmounts resulting from infection with AAV-ShH10Y0CMV-GFP (left column) or AAV-ShH10Y-CMV-Cyp26a1 + AAV-ShH10Y-CMV-GFP (right column). (**J**) Insets of the dorsal peripheral regions in both groups. (**K**) Quantification of CAR^+^ cones in the sampled area of infected retinas (GFP: n=13, Cyp26a1: n=13, Student’s two-tailed T test, *p*=0.0403). ONL, Outer Nuclear Layer; INL, Inner Nuclear Layer; GCL, Ganglion Cell Layer. Scale bars; 500 μm (C top panels, I), 50 μm (F), 100 μm (C bottom panels, J). All results are expressed as the mean*±*SD. *p*<*0.05, **p*<*0.01, ****p*<*0.0001.

Due to the deletion of Aldh1a1, Aldh1a2, and Aldh1a3 in all cells resulting from the use of the broadly active CAG-CreER, it was possible that the effect of peripheral cone death was due to systemic toxicity. For example, hepatocytes in the liver express high levels of Aldh1a1 and regulate lipid metabolism^51^. To overcome this issue, we generated another Aldh Flox mouse line with Pdgfra-Cre, which drives expression of Cre in MG^52^ **(Figure 2F)**. To validate the specificity of the Pdgfra-Cre, the DNA construct, CAG-LoxP-Neo-STOP-LoxP-GFP, which drives GFP expression in Cre^+^ cells was electroporated into the retinas of rd1; Pdgfra-Cre mice at P0. At P10, GFP expression was mostly confined to MG, as evidenced by colocalization with Sox2, a marker of MG and a subset of amacrine cells **(Figure 2F)**. Pdgfra-Cre mice were crossed with Aldh1a1 flox mice to generate a MG-specific Aldh1a1 cKO mouse line. At P60, the retinas and the liver were harvested from littermates with and without Pdgfra-Cre. Using ddPCR, the number of transcripts for *Aldh1a1* was quantified from the retina and the liver. While *Aldh1a1* remained in the liver at high levels, it was essentially depleted in the retinas of mice with Pdgfra-Cre, indicating the specificity of the Cre line (Retina: Mean*±*SD: -CreER: 0.0128*±*0.001; +CreER: 0.0005 *±* 0.0001; Liver: Mean *±* SD: -CreER: 0.5579 *±* 0.6069; +CreER: 0.3255 *±* 0.1793) **(Figure 2G)**. We then quantified the number of CAR^+^ cones in the dorsal retina and found a significant reduction in the number of cones in the retinas with Pdgfra-Cre compared to controls (mean S*±* D: -CreER: 48.62 *±* 10.77; +CreER: 18.73 *±* 7.44) **(Figure 2H)**. These results indicate that MG-specific RA signaling is necessary for peripheral cone survival.

To further investigate whether RA is necessary for peripheral cone survival, another method was used to remove RA from the retina. Overexpression of Cyp26a1, an RA degrading enzyme, in the retina was carried out by AAV transduction. To maximally target MG, we used the ShH10Y capsid, which has a high tropism for retinal MG when injected subretinally or intravitreally^53^. To obtain widespread AAV transduction in MG by subretinal injection, an injection time point was chosen such that most MG would have already been born (*>*P3), and the subretinal space would allow for a wide distribution of the vector. Subretinal injection with AAV-ShH10Y-CMV-GFP into P4 mice achieved high expression of GFP in central and peripheral MG, although some far peripheral MG were not transduced **(Figure 2 – figure supplement 1)**. Subretinal injection into the dorsal retina led to GFP expression throughout the dorsal retina, with partial infection of the ventral retina **(Figure 2I)**. When combined with AAV-ShH10Y-CMV-Cyp26a1, smFISH showed expression of *Cyp26a1* in the dorsal retina **(Figure 2 – figure supplement 2)**. In accord with the cKO data, overexpression of Cyp26a1 significantly reduced the number of dorsal cones compared to GFP only controls (mean *±* SD: GFP: 15.81 *±* 8.369; +CreER: 9.416 *±* 6.564) **(Figure 2J-K)**.

Taken together, these results indicate that RA is necessary for peripheral cone survival in the rd1 mouse model.

### Activation of the RA signaling pathway promotes local cone survival in the central retina

To determine whether activation of the RA signaling pathway is sufficient for cone survival, the signaling pathway was activated in the central retina, independently of RA **(Figure 3A)**. Canonically, the RARa-RXR dimer, bound to genomic RAREs, represses downstream transcription in the absence of the RA ligand^34^ **(Figure 3B)**. In the presence of RA, the dimer recruits co-transcriptional activators and the transcriptional machinery, leading to transcription of downstream genes. The RA signaling pathway is tightly regulated via a type of feedback inhibition, as the major downstream transcriptional target of RAR is Cyp26a1, which breaks down RA. To overcome this issue, a RARa-VP16 transactivator fusion protein was developed previously, to constitutively activate downstream transcriptional targets, even in the absence of RA^54^ **(Figure 3B)**. A plasmid construct encoding this fusion, CAG-RARa-VP16, was electroporated into the ventral-central retina of rd1;RARE-LacZ mice with the electroporation control construct bpCAG-BFP. Retinas were harvested at P40 and cones were quantified in the electroporated patches **(Figure 3A)**. In the bpCAG-BFP only control, the BFP^+^ electroporated patches were localized to the ventral-central retina, but B-gal expression was not induced, and CAR^+^ cones were sparsely distributed **(Figure 3C)**. In contrast, when bpCAG-BFP was combined with CAG-RARa-VP16, a strong induction of B-gal and Cyp26a1 expression was observed, indicative of transcription from RA signaling **(Figure 3C and Figure 3 – figure supplement 1)**. Within the electroporated patches that included CAG-RARa-VP16, a high density of cones was observed **(Figure 3C)**. Quantification of cones in the electroporated region showed a significant increase in the RARa-VP16 group compared to controls (mean *±* SD: BFP: 19.14 *±* 4.589; RARa: 38.8 *±* 6.261) **(Figure 3D and Figure 3 – figure supplement 2)**. Of note, electroporation of the postnatal mouse retina delivers the DNA constructs to late-born cell types of the retina, including rods, MG, bipolar cells, and amacrine cells, but not cones^55^. Therefore, this effect on cone survival is not due to expression of the constructs within cones, i.e. it is a non-cell autonomous effect on cones. Furthermore, the region of increased cone survival was confined to the electroporated patch, with a sharp boundary, suggesting that the effect of RA signaling on cone survival is restricted locally.

**Figure 3:**
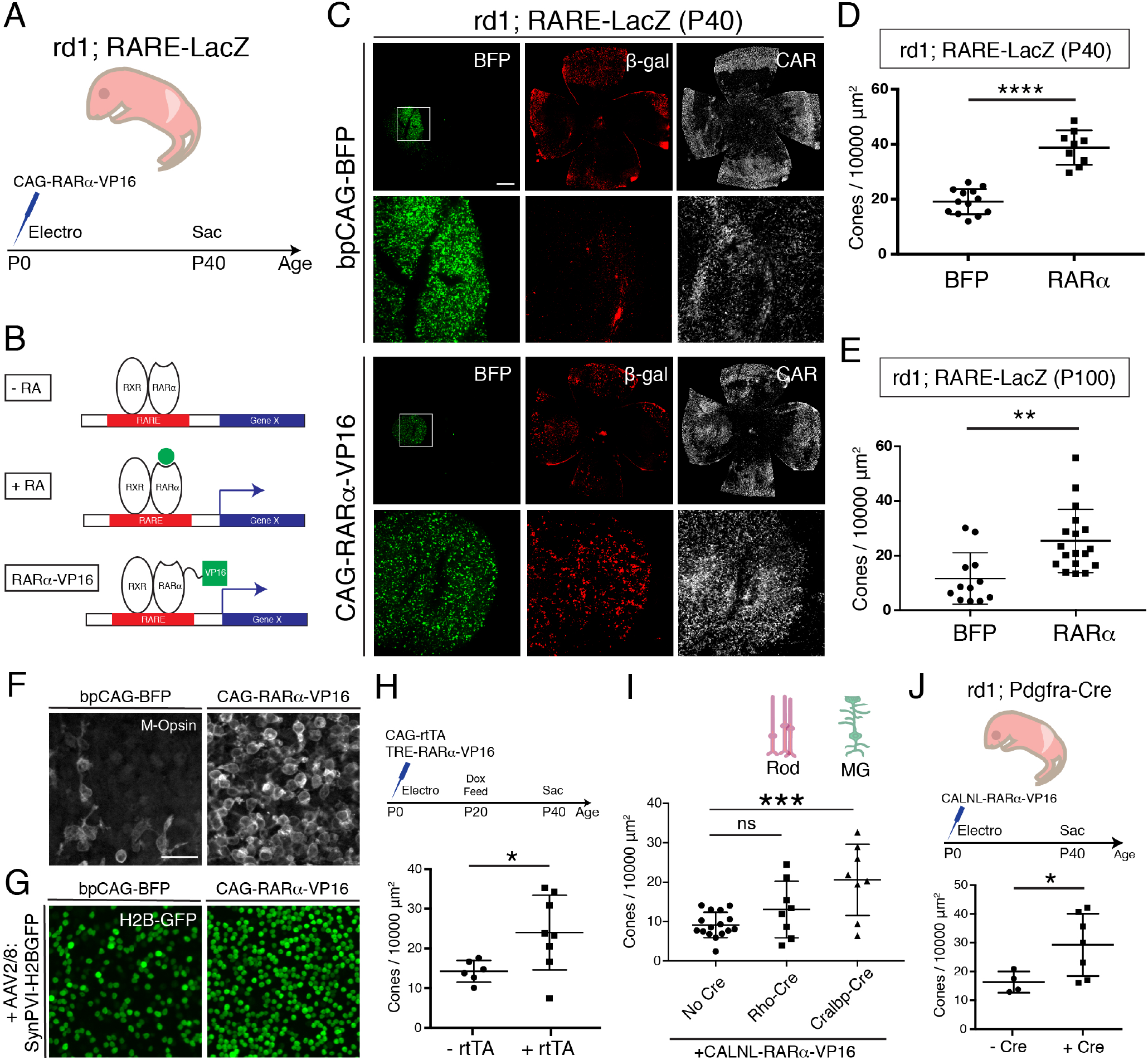
Constitutive RA signaling is sufficient for cone survival in the central retina. (**A**) Schematic of the experimental design. Retinas of rd1; RARE-LacZ mice were electroporated with bpCAG-BFP with or without the CAG-RARa-VP16 construct at P0 or P1 and harvested at P40. (**B**) Schematic of RA-mediated transcriptional activation and RARa-VP16-mediated constitutive activation. Without RA, the RARa-RXR dimer, bound to genomic RAREs, repress downstream transcription. With RA, the dimer activates transcription of downstream genes, which includes the RA degradation enzyme, Cyp26a1, creating a strong negative feedback. With the introduction of RARa-VP16, transcription of genes downstream of RAREs is activated despite the lack of RA. (**C**) IHC against BFP(left column), B-gal (middle column), and CAR (right column) in P40 rd1; RARE-LacZ whole retinal flatmounts (top rows) and BFP^+^ insets (bottom rows) with overexpression of bpCAG-BFP with or without CAG-RARa-VP16. (**D**) Cone quantification in BFP^+^ central retina of P40 rd1; RARE-LacZ mice electroporated with bpCAG-BFP with or without CAG-RARa-VP16 (BFP: n=13, RARa: n=9, Student’s two-tailed T test, p*<*0.0001). (**E**) Cone quantification in BFP^+^ central retina of P100 rd1; RARE-LacZ mice electroporated with bpCAG-BFP with or without CAG-RARa-VP16 (BFP: n=12, RARa: n=18, Student’s two-tailed T test, *p*=0.0019). (**F**) Representative images of IHC against M-Opsin (Opn1mw) in BFP^+^ central retina of P40 rd1; RARE-LacZ mice electroporated with bpCAG-BFP with or without CAG-RARa-VP16. (**G**) Representative images of IHC against GFP in BFP^+^ central retina of P40 rd1; RARE-LacZ mice electroporated with bpCAG-BFP with or without CAG-RARa-VP16 and injected with AAV8-SynPVI-H2BGFP. (**H**) Schematic of the experimental design (top). Retinas of rd1; RARE-LacZ mice were electroporated with TRE-RARaVP16 with or without CAG-rtTA. Doxycycline was administered from P20 – P40, at which time point the retinas were harvested. Cone quantification in BFP^+^ central retina of P40 rd1; RARE-LacZ mice electroporated with with TRE-RARa-VP16 with or without CAG-rtTA (-rtTA: n=6, RARa: n=8, Student’s two-tailed T test, *p*=0.0311). (**I**) Cone quantification in BFP^+^ central retina of P40 rd1; RARE-LacZ mice electroporated with CALNL-RARa-VP16 with or without Rho-Cre or Cralbp-Cre (No Cre: n=16, Rho: n=8, Cralbp: n=8, One-way ANOVA, F=9.372, *p*=0.0007; Tukey’s multiple comparison test, No Cre vs. Rho: *p*=0.3111, No Cre vs. Cralbp: *p*=0.0005, Cralbp vs. Rho: *p*=0.0511). (**J**) Schematic of the experimental design (top). Retinas of rd1; Pdgfra-Cre mice were electroporated with CALNL-RARa-VP16 and harvested at P40. Cone quantification in BFP^+^ central retina of P40 rd1; Pdgfra-Cre electroporated with CALNL-RARa-VP16 (-Cre: n=4, +Cre: n=7, Student’s two-tailed T test, *p*=0.0488). Scale bars; 500 μm (C), 25 μm (F). All results are expressed as the mean *±* SD. *p*<*0.05, **p*<*0.01, ***p*<*0.001, ****p*<*0.0001.

To determine whether the cone survival effect of RA signaling is long lasting, cones were quantified at P100. Based on BFP, the expression level resulting from electroporation diminished over time between P40 and P100. Nevertheless, cone survival was significantly increased in the RARa-VP16 group compared to controls even at P100 (mean*±*SD: BFP: 11.67*±*9.388; RARa: 25.41 *±* 11.58), indicating that this effect lasts over an extended period of time. As RA signaling has been shown to increase the expression level of CAR itself, we sought to verify that RA signaling promoted bona fide cone survival, not just increased CAR expression, as described previously^56^. To this end, IHC for M-opsin (Opn1mw) was carried out, which showed that the surviving cones also expressed M-opsin **(Figure 3F)**. To further verify the identity of these cones, during electroporation, AAV8-SynPVI-H2BGFP was co-delivered. This AAV encodes a construct which drives the expression of nuclear H2B-GFP from a synthetic Gnat2-based cone promoter^57^. The surviving cones in the electroporated patch also expressed H2B-GFP, suggesting that these cones were positive for Gnat2, a cone marker **(Figure 3G)**. These results show that RA signaling leads to survival of cones, instead of simply upregulating the expression of CAR.

Due to the experimental approach of constitutively activating RA signaling by overexpressing CAG-RARa-VP16 via electroporation at P0-P1, it remained unclear which cell type was responsible for the cone survival effect and when the effect was taking place. Electroporation of the retinas of newborn pups leads to expression within a few days in cell types born postnatally, including rods, MG, bipolar cells, and amacrine cells. Therefore, we sought to narrow the time period and cell type(s) that were sufficient for promoting cone survival by RA signaling. To determine the sufficient time point, a Dox-inducible TetON system was used to control the timing of expression. The constructs, CAG-rtTA and TRE-RARa-VP16, were electroporated at P0-P1, and expression was induced from P20 – P40 by Dox administration. With such a paradigm, a significant increase in the number of cones in the +rtTA group compared to controls was observed (mean *±* SD: -rtTA: 14.25 *±* 2.723; +rtTA: 24±9.412), indicating that RA signaling during cone degeneration, and after rod degeneration (after P20), is sufficient for cone survival **(Figure 3H)**. To determine which cell type was involved in promoting cone survival by RA signaling, specific promoters were used to drive expression of RARa-VP16. Rods are the most abundant of the electroporated cell types, while MG are the cell types that expresses Aldh1a1, and thus promoters for these two cell types were used **(Figure 1G)**. Rho-Cre was used to activate expression in rods and Cralbp-Cre was used for expression in MG. Constructs with these promoters were co-electroporated with a Cre-dependent RARa-VP16 construct (CALNL-RARa-VP16). This strategy combined the specificity of the cell type-specific promoters and the high expression level of the CAG promoter. Co-electroporation with Cralbp-Cre, but not Rho-Cre, showed significant increases in the number of cones compared to controls (No Cre) (mean *±* SD: No Cre: 9.125 *±* 3.22; Rho: 13.07 *±* 7.175; Cralbp: 20.61 *±* 9.048) **(Figure 3I)**. To further investigate the cell type question, the CALNL-RARa-VP16 construct was electroporated into rd1; Pdgfra-Cre retinas, which express Cre in MG **(Figure 2F)**. Overexpression of RARa-VP16 in the Cre^+^ retinas increased the number of cones compared to controls without Cre (mean *±* SD: -Cre: 16.32 *±* 3.681; +Cre: 29.25 *±* 10.79) **(Figure 3J)**, in keeping with the results from Cralbp-Cre electroporation.

Taken together, these results indicate that RA signaling activation in MG during cone degeneration (*>*P20) is sufficient to promote cone survival in regions without endogenous RA synthesis.

### *ALDH1A1* is enriched in the peripheral adult human retina

Whether RA signaling plays a role in preserving cones in human RP patients is not known. However, far peripheral islands of vision have been observed in some RP patients by functional visual testing and histology^20,58^. We obtained five postmortem adult human non-RP retinas to investigate the expression level of *ALDH1A1* in the central and peripheral retina. As assessed by ddPCR, *ALDH1A1* was significantly enriched in the peripheral retina **(Figure 4)**. This result opens the possibility that RA signaling may be active in the human peripheral retina and play a role in preserving peripheral vision, if any.

**Figure 4:**
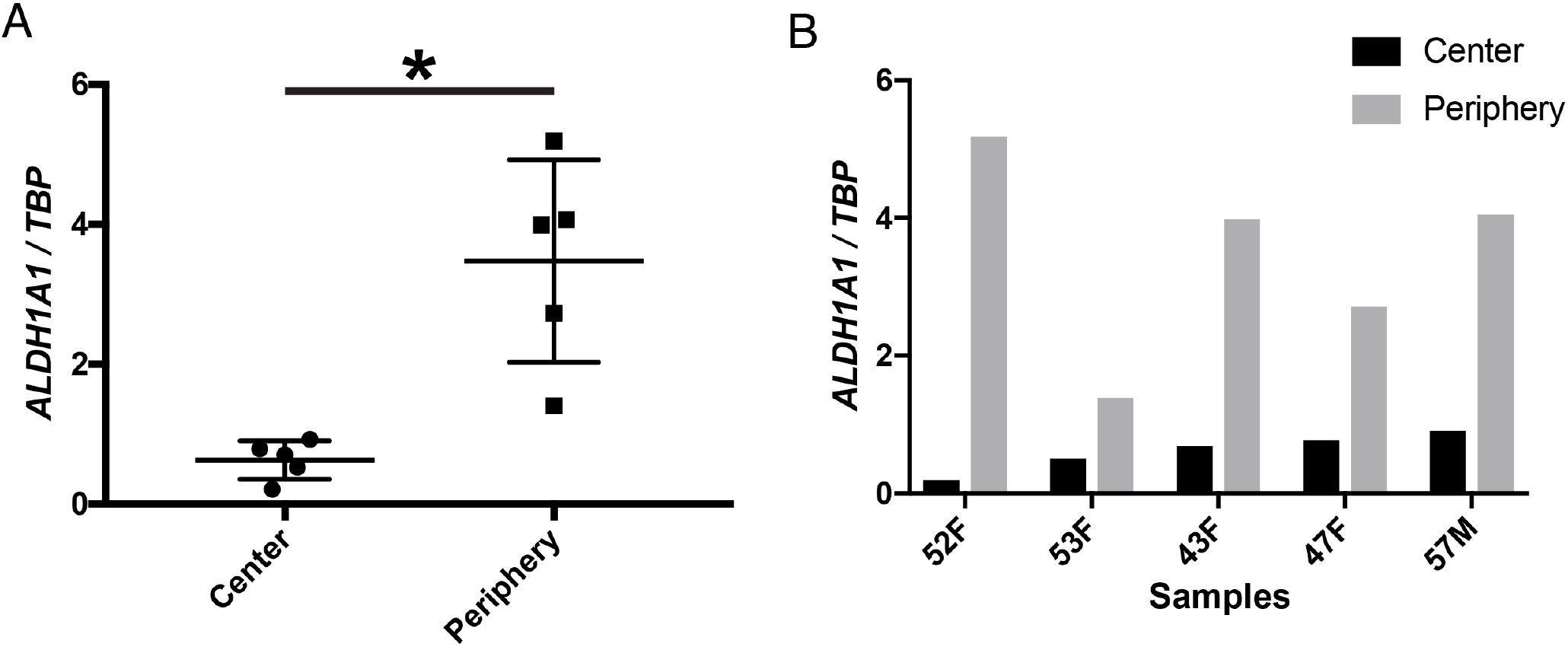
*ALDH1A1* is enriched in the peripheral retina of humans. (**A**) Relative expression level of *ALDH1A1* compared to loading control, TBP, in the center and periphery of the human non-RP retinas (Center: n=5, Periphery: n=5, Two-way ANOVA, *p*=0.0146). (**B**) Expression level of *ALDH1A1* in retinas of individual humans, age and gender as indicated.

## DISCUSSION

In this study, we set out to elucidate the molecular and cellular mechanism underlying long-term peripheral cone survival in the rd1 mouse model of retinal degeneration. We found that peripheral MG express Aldh1a1, leading to active transcription of RA target genes, and that the regions with active RA signaling closely matched those with surviving cones. Removal of RA, either by conditional knockout of the synthesis enzymes or by overexpression of the degradation enzyme, decreased the number of peripheral cones. Conversely, constitutive activation of the RA signaling pathway in MG promoted cone survival in the central retina. Taken together, RA signaling is both necessary and sufficient for cone survival in the rd1 mouse model of retinal degeneration.

We found that peripheral MG express the RA-synthesizing enzyme, Aldh1a1, and activate transcription of RA target genes, autonomously or only within nearby MG. Importantly, RA signaling is not activated directly in cones; and therefore, the mechanism that promotes cone survival is non-cell autonomous with regards to cones. To gain a deeper mechanistic insight into this RA signaling-mediated cone survival, identification of the downstream effector is of great interest. To start, transcriptional profiling of MG and cones from Aldh Flox cKO mice and RARa-VP16 overexpression mice will be helpful to identify candidate ligand-receptor pairs that may play a role in promoting cone survival. However, given the diverse functions of MG, RA signaling may mediate a more complex effect on the tissue environment that secondarily promotes cone survival. For example, RA signaling plays a role in maintaining the integrity of the blood-retina-barrier (BRB) in the zebrafish eye, and BRB leakage is associated with cone degeneration^59,60^. Therefore, it is possible that RA signaling helps keep the BRB in the peripheral retina intact, which in turn helps cones survive, without directly influencing cones. Though the cause of secondary cone degeneration is currently unknown, use of RA signaling as a molecular handle may lead to a deeper mechanistic understanding. For practical applications as a mutation-agnostic gene therapy for patients with RP, identification of the downstream effector will be critical.

To go forward in the clinic, safety and efficacy are key components. For safety, discovery of an effector molecule, which may be less pleiotropic than RARa-VP16 may be important as overexpression of a constitutively active transcription factor will lead to upregulation of myriad genes, potentially with deleterious effects. Gene therapy with an effector molecule also may be more effective than overexpression of RARa-VP16. Currently, AAV-based overexpression of RARa-VP16, even with a strong CMV promoter, was not sufficient to promote cone survival as the expression level was too low. Direct overexpression of an effector molecule(s), with cell type specificity, likely will be a more effective approach for gene therapy. Another reason to identify the downstream effector(s) is to avoid potential retinal ganglion cell (RGC) hyperactivity induced by RA signaling, as was recently reported^61-63^. Although our current gain-of-function and loss-of-function approaches ensure that RGCs are not affected due to the specificity of the promoters and the injection route, it would still be a concern to promote cone survival at the cost of inducing detrimental RGC hyperactivity.

The human central retina is more susceptible to photoreceptor death compared to the peripheral retina in both RP and age-related macular degeneration (AMD)^20,64^, and it remains unclear what molecular mechanisms underlie this difference. One of the markers of the human central retina is the sustained expression of CYP26A1 from embryo into adulthood^65,66^. Furthermore, we have shown that *ALDH1A1* remains enriched in the adult human peripheral retina. While the function of these reciprocal genes in the human retina is unknown, development of the high acuity area in the chick, which lacks rods as does the human fovea, requires inhibition of RA signaling^66^. Though it remains unclear whether CYP26A1 in the human retina is required for foveal development, the lack of RA signaling in the central retina, perhaps as a remnant of development, may play a role in the vulnerability of the central retina to disease such as AMD and RP. Conversely, activated RA signaling by ALDH1A1 may play a role in promoting cone survival in the far peripheral islands of vision seen in patients with RP^20,58^. To further support this hypothesis, it will be beneficial to correlate the expression pattern of ALDH1A1 with regions of extended cone survival in people with RP.

## MATERIALS AND METHODS

### Mouse

All animals were handled according to protocols approved by the Institutional Animal Care and Use Committee (IACUC) of Harvard University (IACUC protocol: 1695). RARE-LacZ mice (stock #008477), FVB/rd1 mice (stock #207), rd10 mice (stock #004297), CAG-CreER mice (stock #004682), and Pdgfra-Cre mice (stock #013148) were obtained from Jackson Laboratory. Rho^-/-^mice were a gift from Janis Lem, Tufts University, Boston, MA^67^. Aldh1a1, 2, 3 flox/flox mice were a gift from Norbert Ghyselinck, IGBMC, France. All retina degeneration mouse lines were housed within +/-2 rows in their respective racks to control for light exposure. For tissue harvest, mice were euthanized with CO_2_ and then secondarily with cervical dislocation. All genotyping primer sequences are in Supplementary Files.

### Human Retina Samples

Frozen eyes were obtained from Restore Life USA (Elizabethton, TN) through TissueForResearch. Patient DRLU032618A was a 52-year-old female with no clinical eye diagnosis and the postmortem interval was 8 hours. Patient DRLU041518A was a 57-year-old male with no clinical eye diagnosis and the postmortem interval was 5 hours. Patient DRLU041818C was a 53-year-old female with no clinical eye diagnosis and the postmortem interval was 9 hours. Patient DRLU051918A was a 43-year-old female with no clinical eye diagnosis and the postmortem interval was 5 hours. This IRB protocol (IRB17-1781) was determined to be not human subjects research by the Harvard University-Area Committee on the Use of Human Subjects.

### Plasmids

Rho-Cre and Cralbp-Cre were from Matsuda and Cepko^68^. Newly generated plasmids (CAG-RARa-VP16, CALNL-RARa-VP16, TRE-RARa-VP16, CAG-rtTA, AAV-CMV-Cyp26a1) have been deposited to Addgene.

### Retina Flatmount IHC

Eyes were enucleated, and the retinas (with the lens) were dissected in PBS. These retinas were fixed in 4% PFA (Electron Microscopy Sciences, cat. #15714S, diluted in PBS) for 20 minutes at RT. After 2x washes with PBS, they were incubated in Blocking Buffer (0.3% Bovine Serum Albumin (Jackson ImmunoResearch, cat. #001-000-162), 4% donkey serum (Jackson ImmunoResearch, cat. #017-000-121), 0.3% Triton X-100 (Sigma Millipore, cat. #T8787), diluted in PBS) for 15 minutes at RT. Retinas were incubated in primary antibody, diluted in Blocking Buffer, for 2 -4 hours at RT with the following antibodies at the following concentrations: rabbit Cone Arrestin (1:3000, Millipore Sigma, cat. #AB15282), chicken B-galactosidase (1:3000, Aves Labs, cat. #BGL1010), goat B-galactosidase (1:3000, AbD Serotec, cat. #4600-1409), mouse Brn3a (1:500, Millipore Sigma, cat. #MAB1585), rabbit Iba1 (1:500, GeneTex, cat. #GTX100042), mouse Glutamine Synthetase (1:3000, Millipore Sigma, cat. #MAB302), goat Choline Acetyltransferase (1:500, Millipore Sigma, cat. #AB144P), chicken GFP (Aves Labs, cat. #GFP-1020), mouse Sox2 (1:500, RD Systems, cat. AF2018), rabbit Opn1mw (Millipore Sigma, cat. #AB5405), goat Aldh1a1 (Abcam, cat. #9883, 0.5% Triton X-100 was used in the Blocking Buffer for this antibody). After 3x wash with PBS, the retinas were incubated in corresponding secondary antibodies conjugated to AlexaFluor 488, 594, or 647, diluted in Blocking Buffer (1:750, Jackson ImmunoResearch) for 1 hour at RT. After 3x washes with PBS, the lens were removed, and the retinas were flatmounted on a No. 1.5 coverslip (VWR, cat. #48393-241) and dried. The coverslips were mounted on a Superfrost Plus slides (Fisher Scientific, cat. #12-550-15) with Fluoromount-G (SouthernBiotech cat. #0100-01) and dried overnight at RT.

### Retinal Flatmount LacZ Staining

LacZ staining on RARE-LacZ flatmount was performed as described previously^66^.

### Retina Cryosectioning

Eyes were enucleated, and the retinas (with the lens) were dissected in PBS. The retinas were fixed in 4% PFA for 20 minutes at RT. For smFISH experiments, it was critical to use freshly opened PFA ampule for fixation. After 2x washes with PBS, the lens were removed. For experiments that required only the electroporated region, the BFP^+^ area was dissected out under a fluorescent dissection scope. The retina was incubated in 30% sucrose/PBS (Millipore Sigma, cat. #S0389) until the tissue sank to the bottom. The tissue was embedded in 50%/15% OCT/Sucrose solution (100% OCT mixed with 30% sucrose, equal parts, Tissue-Tek, cat. #25608-930) in a cryomold. The retinas were snap-frozen in dry ice/ethanol slurry and kept at -80^°^C. The retinas were cryosectioned at 30 μm thickness and mounted onto Superfrost plus slides. The slides were stored at -80^°^C.

### Retina Section IHC

Slides with retina sections were 2x washed with 3 mL of PBS and then dried completely. The slides were washed 1x with PBS. They were incubated with 500 μL of Blocking Buffer for 15 minutes at RT and then with 500 μL of primary antibody diluted in Blocking Buffer for 2 hours at RT. After 3x washes with PBS, the slides were incubated with 500 μL of secondary antibody diluted in Blocking Buffer for 1 hour at RT. After 3x washes with PBS, the slides were dried completely and coverslipped with Fluoromount-G.

### Single Molecule FISH

SABER FISH and RNAscope were used for smFISH of retinal sections^69^. For SABER, the slides were washed with PBS for 5 - 10 minutes to remove the OCT on the slides. Subsequently, sections were completely dried, and an adhesive chamber (Grace Bio-Labs, cat. #621502) was placed to encase the sections. The samples were incubated in 0.1% PBS/Tween-20 (MilliporeSigma, cat. #P9416) for 10 minutes. The PBST was removed, and the samples were incubated with pre-warmed (43^°^C) 40% wHyb (2x SSC (Thermo Fisher Scientific, cat. #15557044), 40% deionized formamide (MilliporeSigma, cat. #S4117), 1% Tween-20, diluted in UltraPure Water) for at least 15 minutes at 43^°^C. The 40% wHyb was removed, and the samples were incubated with 100 μL of pre-warmed (43^°^C) Probe Mix (1 μg of probe per gene, 96 μL of Hyb1 solution (2.5x SSC, 50% deionized formamide, 12.5% Dextran Sulfate (MilliporeSigma cat. #D8906), 1.25% Tween-20), diluted up to 120 μL with UltraPure Water) and incubated 16 – 48 hours at 43^°^C. The samples were washed twice with 40% wHyb (30 minutes/wash, 43^°^C), twice with 2x SSC (15 minutes/wash, 43^°^C), and twice with 0.1% PBST (5 minutes/wash, 37^°^C). The samples were then incubated with 100 μL of Fluorescent Oligonucleotide Mix (100 μL of PBST, 2 μL of each 10 μM Fluorescent Oligonucleotide) for 15 minutes at 37^°^C. The samples were washed three times with PBST at 37^°^C for 5 minutes each and counterstained with DAPI (Thermo Fisher Scientific, cat. #D1306; 1:50,000 of 5 mg/mL stock solution in PBS). The oligo sequences for the probes (*Aqp4, Aldh1a1, Cyp26a1*) are in Supplementary Files. For RNAscope, the RNAscope Fluorescent Multiplex Assay was used with 30 μm cryosections according to protocol (Advanced Cell Diagnostics). The following probes were used: Mm-Arr3 (cat. #486551), Mm-Nrl (cat. #475011).

### *in vivo* Electroporation

Electroporation of DNA plasmids into neonatal mouse retina was performed as described previously^55,70^. Briefly, glass needles were created by pulling Wiretrol II capillaries (Drummond Scientific Company, cat. #5-000-2005) using a needle puller (Sutter Instrument, Model P-97). The glass needles were beveled on two edges with a microgrinder (Narishige, cat. #EG-401). 10 μL of DNA plasmids (1 μg/μL per construct) was prepared with 0.5 μL of 2.5% FastGreen (Millipore Sigma cat. #F7252). P0 – P1 mouse pups were anesthetized by cryoanesthesia on ice. A small skin incision was made by a 30-gauge needle and the eye was exposed. The beveled glass needle containing the DNA plasmid was inserted through the sclera into the subretinal space, and the DNA solution was injected into the ventral hemisphere using a Femtojet Express pressure injector (Eppendorf, cat. #920010521) with the following setting: 330 Pa, 3 seconds. The eyelids were then closed with a cotton applicator. Tweezer-type electrodes (Harvard Apparatus, BTX, model 520, 7mm diameter, cat. #450165) were placed with the positive end slightly above the injected eyelid. An electric field was applied using an electroporator (BEX, cat. #CUY21EDIT) with the following parameters: Volts: 80 V, Pulse-On: 50 ms, Pulse-Off: 950 ms, Number of pulses: 5. Then, the eyelids were dried with a cotton applicator to ensure eyelid closure.

### AAV Production

HEK293T cells were seeded onto five 15 cm plates (Celltreat cat. #229651) and grown in 10% FBS/DMEM/PS media (Thermo Fisher Scientific, cat. #10437028, cat. #11995065, cat. #15140163). At 100% confluency, the cells were transfected using 340 μL of polyethylenimine (PolyScience, cat. #24765-2, diluted to 1 mg/mL) with the following DNA plasmids: 35 μg of ShH10Y capsid (a gift from John Flannery, Berkeley, CA), 35 μg of AAV-CMV-Cyp26a1 or AAV-CMV-GFP vector, 100 μg of pHGTI-adeno1. The cells were incubated in 10% NuSerum/DMEM/PS (BD Biosciences, cat. #355500) for 24 hours. Then, the media was changed to DMEM without serum for 48 hours. Next, the cells and media were collected and centrifuged at 1000 xg for 5 minutes at RT. The cell pellet (1) and the media (2) were separated for downstream processing. (1) The cell pellet was resuspended in 10 mL of lysis buffer (150 mM NaCl, 20 mM Tris-HCl pH 8.0) and underwent 3x freeze-thaw between dry ice/ethanol bath and 37^°^C water bath. 10 μL of 1M MgCl_2_ and 10 μL of Benzonase (Millipore Sigma, cat. #E1014) were added and incubated at 37^°^C for 15 minutes. It was then centrifuged at 3800 xg for 20 minutes at 4^°^C, and the supernatant was collected. (2) The media was filtered through a 0.45 μm CA filter (Corning, cat. #430768). NaCl (to 0.4M) and PEG8000 (to 8.5% w/v, Sigma, cat. #81268) were added over a period of 1.5 hours with stirring at 4^°^C. It was then centrifuged at 7000 xg for 10 minutes at 4^°^C and the supernatant was discarded. The supernatant from (1) was added and the viral pellet was resuspended. This solution was overlaid on top of a gradient (17%, 25%, 40%, 60%) of Iodixanol (Millipore Sigma, cat. #D1556) and ultracentrifuged at 46,500 rpm (Beckman Coulter VTi50) for 1.5 hours at 16^°^C. The viral fraction in the 40% gradient was harvested with a 21-gauge needle, and PBS was added up to 15 mL. The solution was transferred to an Amicon tube with a centrifugal filter (Millpore Sigma, cat. #UFC910024) and centrifuged at 1500 xg for 15 minutes at 4^°^C. This process was repeated three times with fresh PBS. At the end of the PBS washes, the remaining viral volume (approximately 100 μL) was collected and titered based on the intensity of VP1, VP2, and VP3 proteins on an SDS-PAGE gel.

### AAV injection

All AAV injections were performed at P4. Subretinal injections were performed as described above (under in vivo Electroporation). Per eye, approximately 5E8 vector genomes (vg) were delivered into the dorsal subretinal space.

### Doxycycline Administration

At P20, mice electroporated with or without CAG-rtTA and TRE-RARa-VP16 constructs were fed Doxycycline-added diet (2000 mg/kg, Envigo, cat. #TD.05512) ad libitum. The diet was stored at 4^°^C and replaced once per week.

### Tamoxifen injection

Aldh Flox mice were intraperitoneally injected with tamoxifen, diluted in corn oil (20 mg/mL, 100 μL per pup, Millipore Sigma, cat. #T5648) at P10. To dissolve the tamoxifen in corn oil (Millipore Sigma, cat. #C8267), it was heated to 65^°^C in a water bath for 1 – 2 hours with occasional vortexing. The dissolved tamoxifen was aliquoted and stored at 4^°^C for less than 1 week.

### Statistical Analysis

Student’s two-tailed T test, one-way ANOVA with Tukey’s multiple comparison test, and two-way ANOVA were performed to compare between control and experimental groups. Multiple litters were used for each experiment, but each litter contained both control and experimental groups for all experiments to account for litter-to-litter variability. Both sexes were used randomly.

### ddPCR

Retina or liver tissue was harvested into 500 μL of Trizol (Thermo Fisher Scientific, cat. #15596026). The tissue was homogenized with disposable pestles (Fisher Scientific, cat. #12141368), and RNA was extracted according to the Trizol protocol. cDNA was generated using either the Superscript 3 or 4 First Strand Synthesis System (Thermo Fisher Scientific, cat. #18080051 or #18091050). For ddPCR, QX200 EvaGreen 2x Supermix was used (BioRad, cat. #1864034) with 0.5 μm of primers and appropriate amounts of cDNA. The primer sequences can be found in Supplementary Files.

### Cell type specific bulk RNA sequencing

Neural retinas (without RPE) from P28 – P36 CD1 WT mice were dissected in PBS and flatmounted. The far peripheral 1/3 of each flatmount petal, each comprising roughly one quarter of each retina, was collected as peripheral retina tissue. The central 1/3 was collected as central retina tissue. The mid-peripheral 1/3 was discarded. The retina was then transferred to a microcentrifuge tube and incubated for 7 minutes at 37°C with an activated papain dissociation solution (87.5 mM HEPES pH 7.0 (Thermo Fisher Scientific, cat. #15630080), 2.5 mM L-Cysteine (MilliporeSigma, cat. #168149), 0.5 mM EDTA pH 8.0 (Thermo Fisher Scientific, cat. #AM9260G), 10 μL Papain Suspension (Worthington, cat. #LS0003126), 19.6 μL UltraPure Nuclease-Free Water (Thermo Fisher Scientific, cat. #10977023), HBSS up to 400 μL, activated by a 15-minute incubation at 37°C). The retina was then centrifuged at 600 xg for 3 minutes. The supernatant was removed, and 1 mL of HBSS/10% FBS (Thermo Fisher Scientific, cat. #10437028) was added. The pellet was then centrifuged at 600 xg for 3 minutes. The supernatant was removed, and 600 μL of trituration buffer (DMEM (Thermo Fisher Scientific, cat. #11995065), 0.4% (wt/vol) Bovine Serum Albumin (MilliporeSigma cat. #A9418)) was added. The pellet was dissociated by trituration at room temperature (RT) using a P1000 pipette up to 20 times or until the solution was homogenous. For all remaining solutions, 5 μL/mL RNasin Plus (Promega, cat. #N2611) was added 10 minutes before use. The cells were centrifuged at 600 xg for 3 minutes, and the pellet was resuspended in 1 mL of 4% PFA and 0.1% saponin (Millipore Sigma, cat. #47036) for 30 minutes at 4^°^C. After 2x wash in Wash Buffer (0.1% saponin, 0.2% BSA), the cells were incubated in primary antibody (Cone Arrestin, 1:1000, in 0.1% saponin/1% BSA) for 30 minutes at 4^°^C. The cells were washed twice in Wash Buffer and incubated in secondary antibody (Donkey anti-rabbit 647 in 0.1% saponin/1% BSA) for 30 minutes at 4^°^C. After 1x wash, the sample was passed through a 35 μm filter (Thermo Fisher Scientific, cat. #352235) before proceeding to FACS. FACSAria (BD Biosciences) with 488, 561, 594, and 633 nm lasers was used for FACS. CAR^+^ population was sorted into microcentrifuge tubes with 500 μL of PBS and kept on ice after FACS. The sorted cells were transferred to a 5-mL polypropylene tube and centrifuged at 3000 xg for 7 minutes at RT. The supernatant was removed, and the cells were resuspended in 100 μL Digestion Mix (RecoverAll Total Nuclear Isolation Kit (Thermo Fisher Scientific, cat. #AM1975) 100 μL of Digestion Buffer, 4 μL of protease), and incubated for 3 hours at 50^°^C, which differs from the manufacturer’s protocol. The libraries for RNA sequencing were generated using SMART-Seq v.4 Ultra Low Input RNA (Takara Bio, cat. #634890) and Nextera XT DNA Library Prep Kit (Illumina, cat. #FC1311096) according to the manufacturer’s protocol. 11 cycles were used for the SMART-Seq. The cDNA library fragment size was determined by the BioAnalyzer 2100 HS DNA Assay (Agilent, cat. #50674626). The libraries were sequenced as 75bp paired-end reads on NextSeq 500 (Illumina). The RNA-seq data was analyzed as described previously_71_.

### Image Acquisition

Fluorescent flatmount images were acquired with Nikon Ti inverted widefield microscope with a Prior ProScanIII motorized stage. The objective used was Plan Apo Lambda 10x/0.45 Air DIC N1 objective, and the camera used was Hamamatsu ORCA-Flash 4.0 V3 Digital CMOS camera. Fluorescent retina section images were acquired with W1 Yokogawa Spinning disk confocal microscope with 50 μm pinhole disk and 488, 561, and 640 nm laser lines. The objectives used were either Plan Apo 20x/0.75 air or Plan Apo 60x/1.4 oil objectives, and the camera used was Andor Zyla 4.2 Plus sCMOS monochrome camera. Nikon Elements Acquisition Software (AR 5.02) was used for image acquisition and Fiji or Adobe Photoshop CS6 was used for image analysis.

### Cone Quantification

For electroporated retinas, the number of cones were quantified in central BFP^+^ regions as schematized in Figure 3 – figure supplement 2. Briefly, a circle of 1.5mm in radius was drawn with the center overlaid on the optic nerve head. This method was used to ensure that the far peripheral cones were not quantified. Within this circle, the BFP^+^ region was used for counting CAR^+^ cones. For unelectroporated retinas, a 640 μm x 640 μm square box was drawn in the middle of the dorsal region, 70% of the length from the optic nerve head, as schematized in Figure 2D. For all quantifications, cones were counted blinded using the Cell Counter function in Fiji, and in some instances, verified by a second blinded counter. For all comparisons between conditions, both conditions were included among littermates to account for litter-to-litter variability. The area of electroporated region was calculated using Fiji.

## Data availability

Raw sequencing data and matrices of read counts are available at GEO: GSE186612.

## Acknowledgments

We would like to thank former and current members of the Cepko and Tabin Labs for the insightful discussions and feedback. We thank P.M. Llopis, R. Stephansky, and the MicRoN core at Harvard Medical School for their assistance with microscopy. We thank C. Araneo, F. Lopez, and the Flow Cytometry Core Facility for their assistance with flow cytometry. We thank J. Patrice and the HCCM animal facility for their assistance with mouse husbandry. This work was supported by the Howard Hughes Medical Institute (C.L.C.), NIH K99/R00 Pathway to Independence Award (R.A. K99EY032110), and Edward R. and Anne G. Lefler Postdoctoral Fellowship (R.A.).

## Author Contributions

R.A. and C.L.C. conceived the study and designed the experiments. R.A. executed the experiments and analyzed the data. G.W. assisted with various experiments. R.A. and C.L.C. wrote the manuscript, and all authors edited the manuscript. C.L.C. supervised all aspects of the work.

