## Supplementary material for "Retinoic acid signaling mediates peripheral cone photoreceptor survival in a mouse model of retina degeneration": Figure Supplements

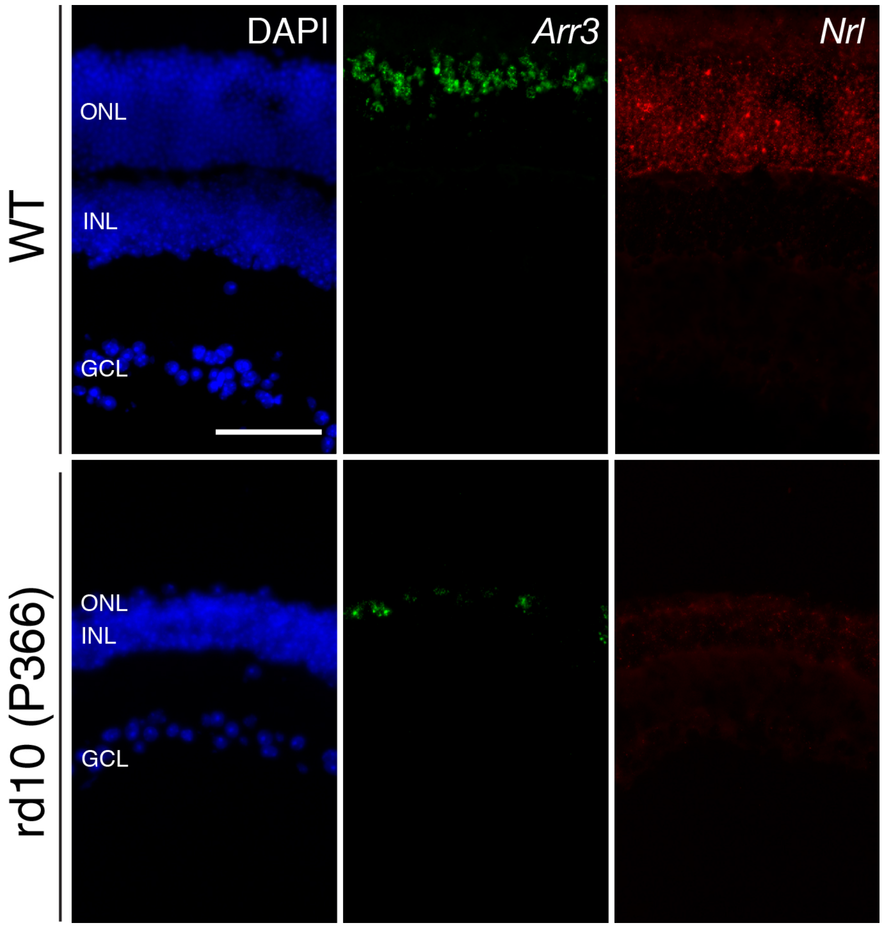


**Figure 1 – figure supplement 1: Long-term cone survival despite complete rod loss.**

Representative images of smFISH against *Arr3* (middle panels) and *Nrl* (right panels), cone and rod markers, respectively, in WT (top panels) and P366 rd10 (bottom panels) retinas, counterstained with DAPI. Scale bar; 50 μm.

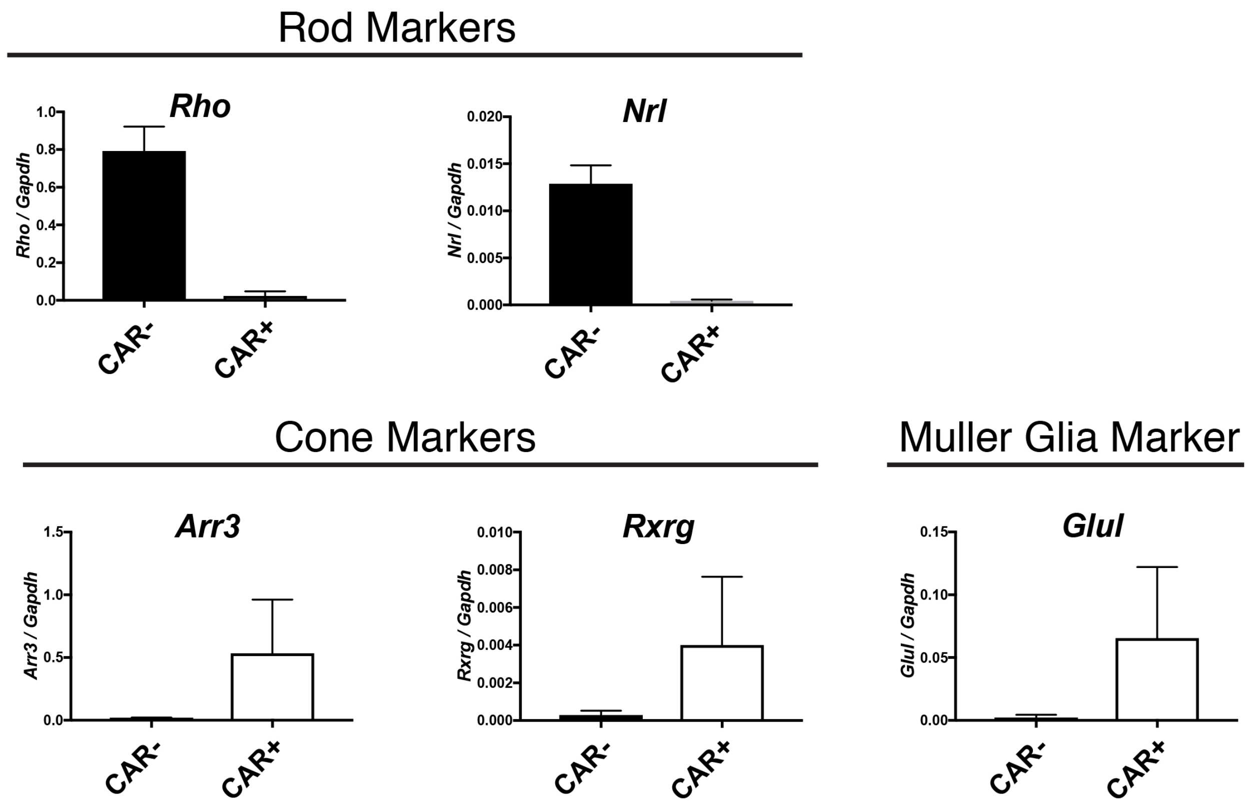


**Figure 1 – figure supplement 2: Cone and MG enrichment in CAR$^+$ FACS-isolated cells.**

Relative expression of rod (*Rho, Nrl)*, cone (*Arr3, Rxrg*), and MG (*Glul)* markers in CAR$^+$ and CAR$^-$ FACS-isolated populations from CD1 mice as detected by ddPCR.

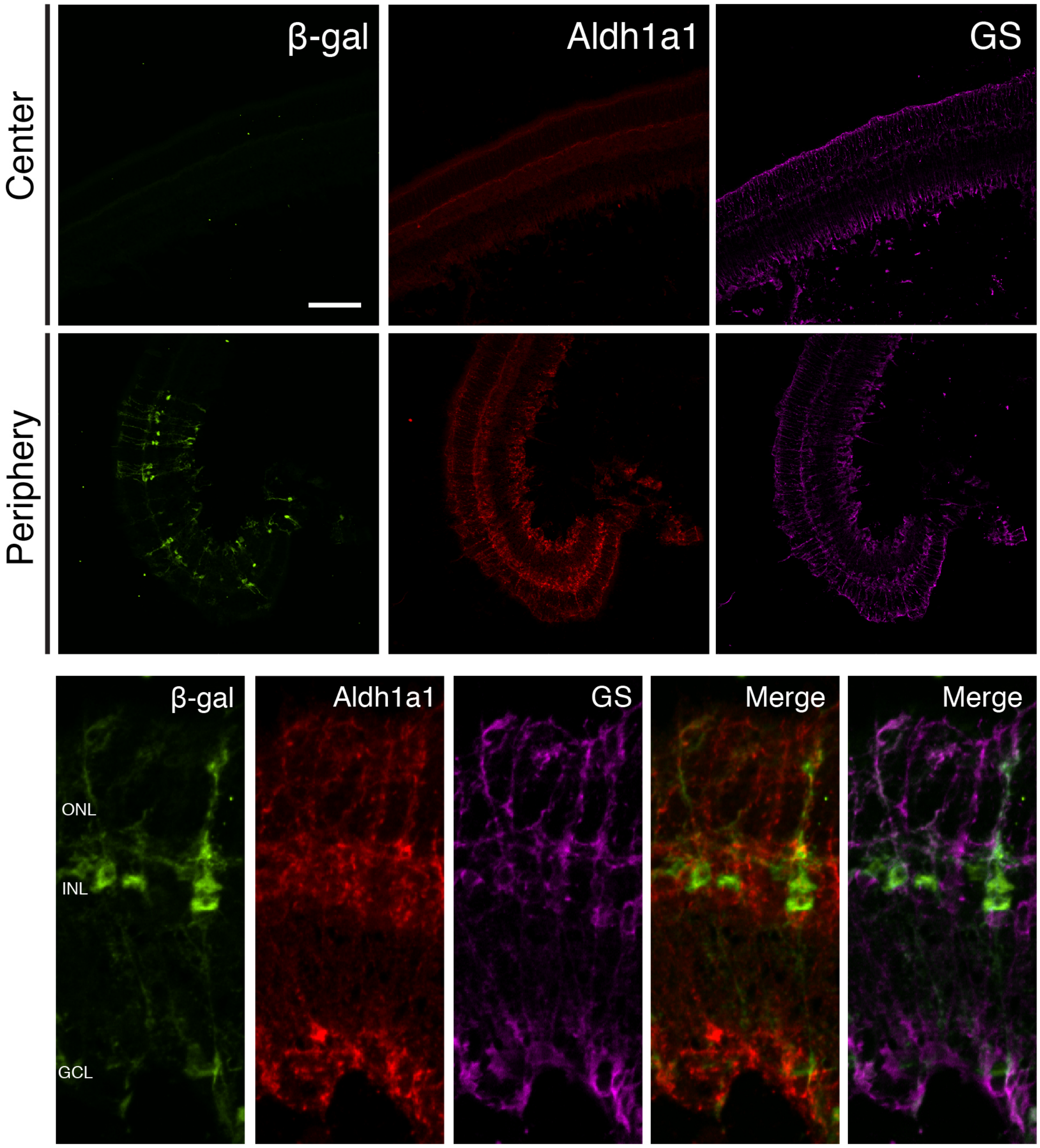


**Figure 1 – figure supplement 3: Aldh1a1 is expressed and RA signaling is active in peripheral MG.**

IHC against B-gal, Aldh1a1, and GS in adult RARE-LacZ (non-degenerating) retinas showing the central (top panels) and peripheral (bottom panels) regions. Scale bar; 100 µm.

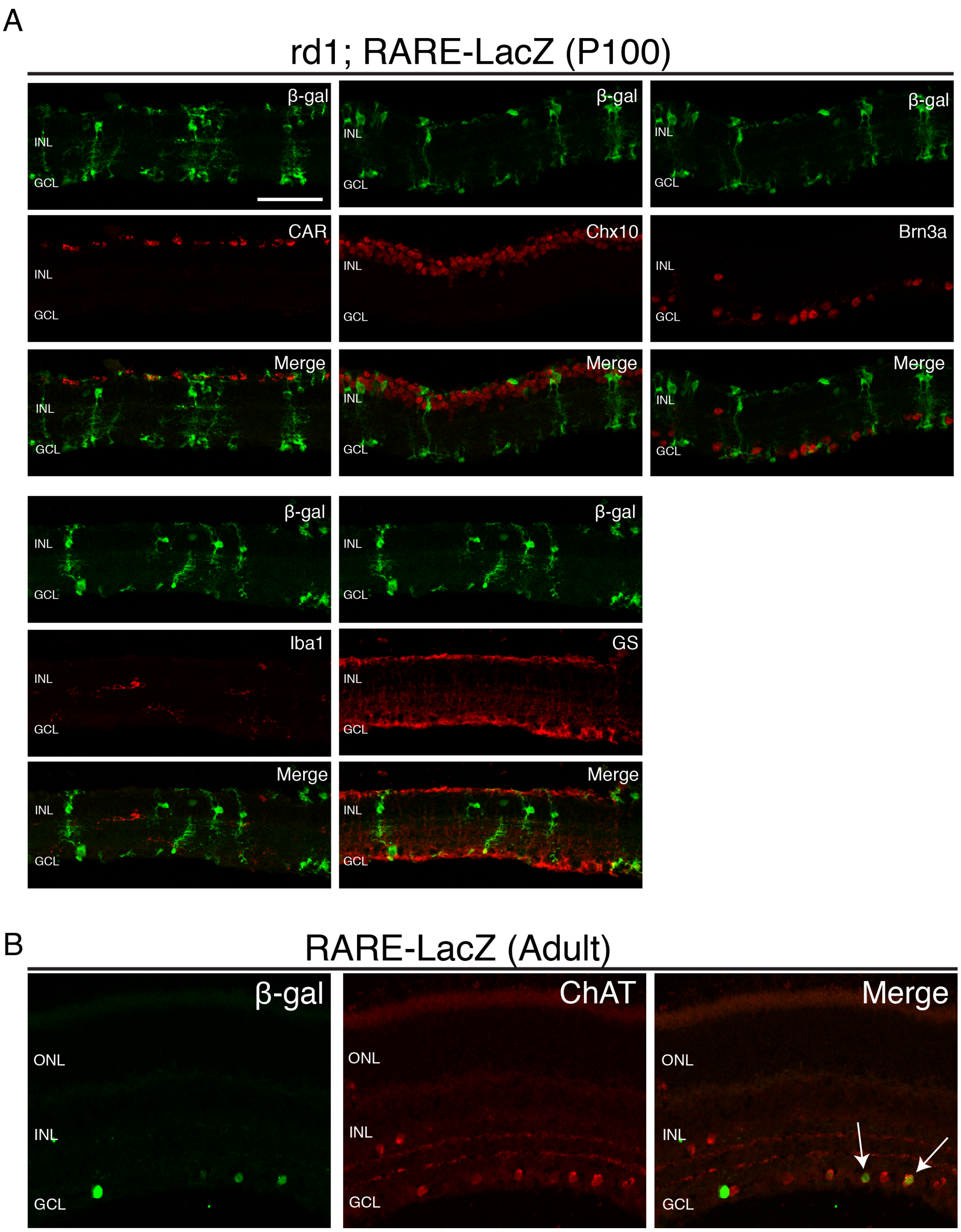


**Figure 1 – figure supplement 4: RA signaling activity is detected only in MG and a subset of ChAT$^{+}$ amacrine cells.**

IHC against B-gal and retina cell type specific markers, including CAR (cones), Chx10 (bipolar cells and MG), Brn3a (RGCs), Iba1 (microglia), GS (MG), and ChAT (amacrine cells) in P100 rd1; RARE-LacZ retinas. Scale bar; 50 μm.

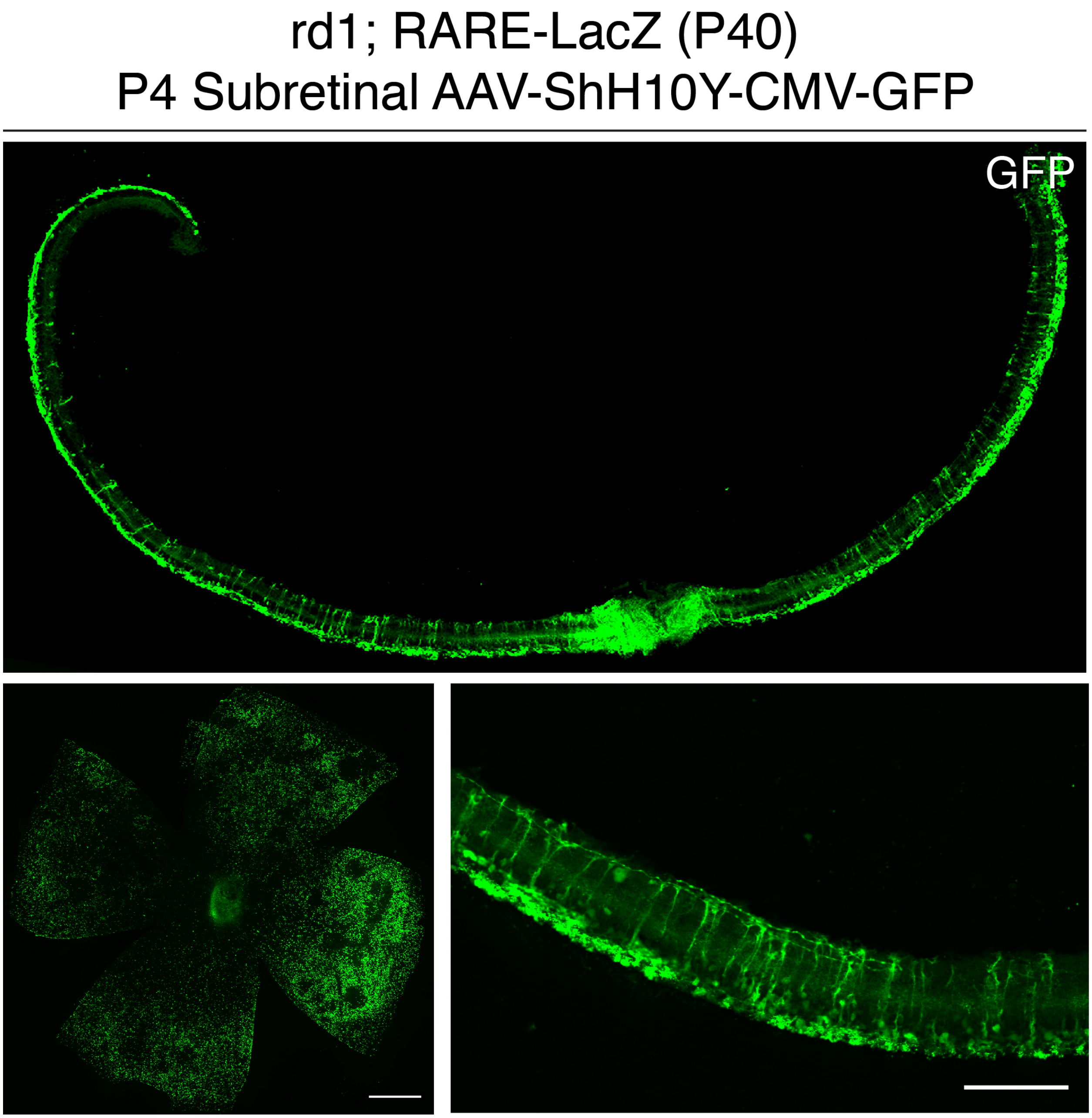


**Figure 2 – figure supplement 1: P4 subretinal injection of AAV-ShH10Y-CMV-GFP leads to retina-wide infection of MG and photoreceptors.**

IHC against GFP in sections and flatmounts of P40 rd1; RARE-LacZ retinas infected with AAV-ShH10Y-CMV-GFP after P4 subretinal injection. Scale bars; 500 µm (left panel), 100 µm (right panel).

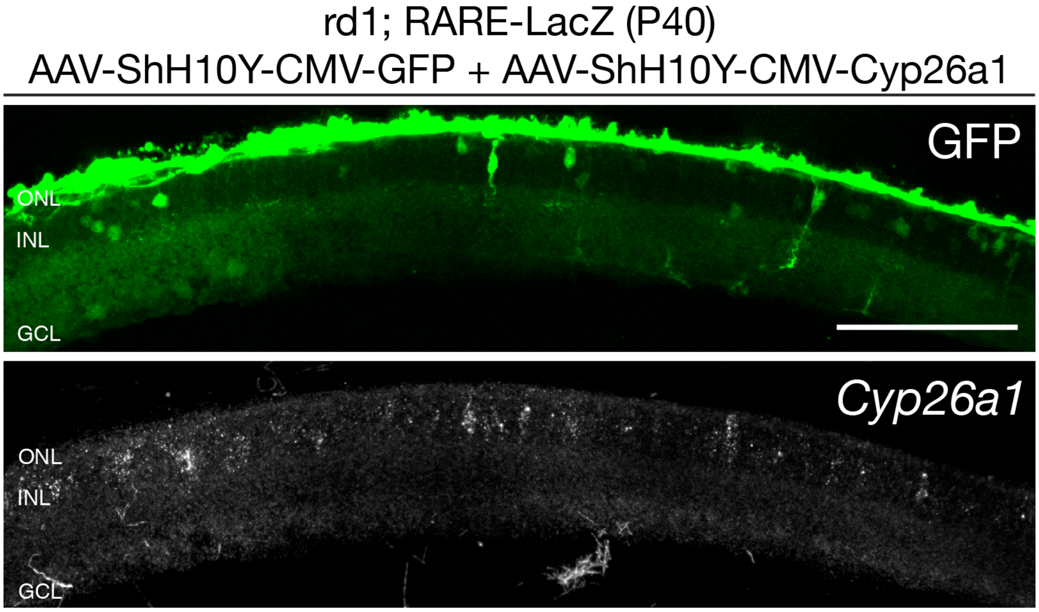


**Figure 2 – figure supplement 2: *Cyp26a1* is overexpressed upon infection with AAV-ShH10Y-CMV-Cyp26a1.**

smFISH against *Cyp26a1* using SABER FISH in P40 rd1; RARE-LacZ retina sections infected with AAV-ShH10Y-CMV-Cyp26a1 and AAV-ShH10Y-CMV-GFP. Scale bar; 100 µm.

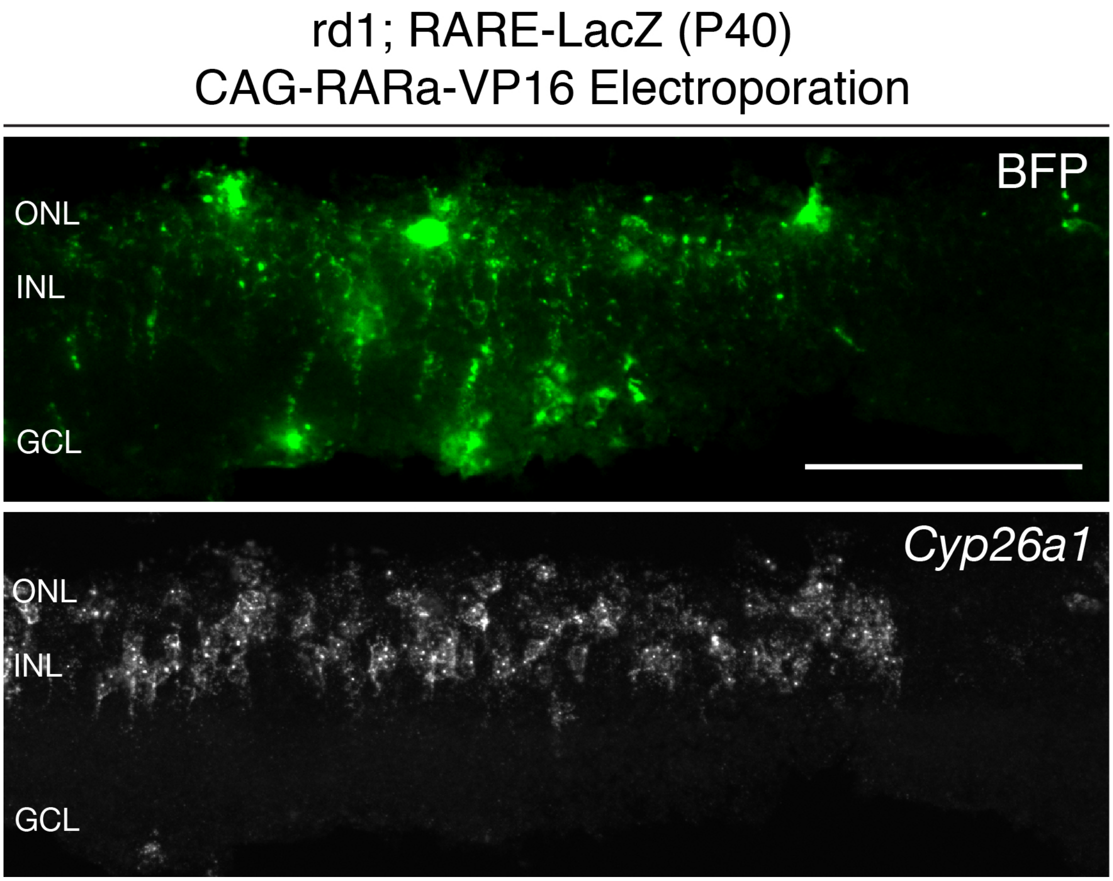


**Figure 3 – figure supplement 1: *Cyp26a1* expression is upregulated upon RARa-VP16 overexpression.**

smFISH against *BFP* and *Cyp26a1* using SABER in P40 rd1; RARE-LacZ retina sections electroporated with CAG-RARa-VP16. Scale bar; 100 μm.

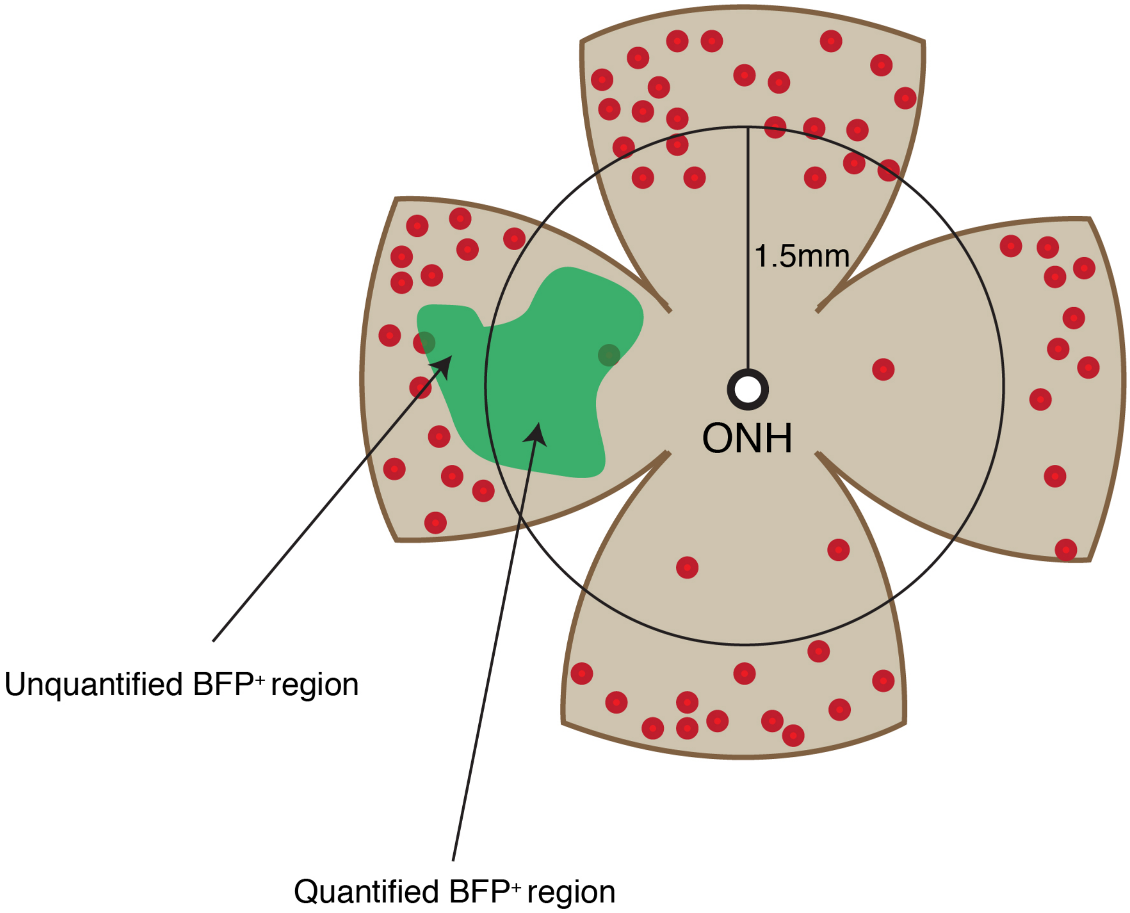


**Figure 3 – figure supplement 2: Cone quantification strategy in gain-of-function experiments.**

Schematic of the area of cone quantification in the dorsal retina. Red dots represent surviving cones, and the green region is the area of electroporation. A circle with a radius of 1.5 mm was used to crop the central retina to ensure that the naturally surviving peripheral cones were excluded. Only the cones in the central BFP$^+$ region were counted.
